# Alternative paths to immune activation: the role of costimulatory risk genes for polygenic inflammatory disease in T helper cells

**DOI:** 10.1101/2022.11.23.517727

**Authors:** Alexandru-Ioan Voda, Kristina Correa, Jonathan Hamp, Chloe Moscrop, Michael Dustin, Luke Jostins-Dean

**Author notes:** These authors contributed equally (Alexandru-Ioan Voda, Kristina Correa, Jonathan Hamp).

## Abstract

T cell activation pathways have been repeatedly implicated by genetic studies as being enriched for risk genes for immune and inflammatory diseases. Many of these risk genes code for costimulatory receptors or ligands. Costimulatory receptors are cell surface proteins on T cells, which are engaged by costimulatory ligands on antigen-presenting cells. Both costimulation and antigen binding are required to trigger T cell activation. In order to study the different pathways activated by these costimulatory risk molecules, and the role they may play in inflammatory disease genetics, we carried out gene expression (RNA-seq) and chromatin accessibility (ATAC-seq) profiling of naive and memory CD4+ T cells (N=5 donors) activated via four different costimulatory receptors: CD28 (the standard molecule used for *in vitro* activation studies), along with alternative costimulatory molecules ICOS, CD6, and CD27.

Most, but not all, activation genes and regions are shared by different costimulation conditions. Alternative costimulation induced lower proliferation and cytokine production, but higher lysosome production, altered metabolic processing, and indications of “signal seeking” behaviour (homing and expression of costimulatory and cytokine receptors). We validated a number of these functions at the surface protein level using orthogonal experimental techniques. We found the strongest enrichment of heritability for inflammatory bowel disease in shared regions upregulated by all costimulatory molecules. However, some risk variants and genes were only induced by alternative costimulation, and the impact of these variants on expression were less often successfully mapped in studies of T cells activated by traditional CD28 costimulation. This suggests that future genetics studies of gene expression in activated T cells may benefit from including alternative costimulation conditions.

## Introduction

Genetic studies of immune and inflammatory diseases have identified many hundreds of risk variants and genes across a wide range of pathways. Cellular functions of CD4+ T cells have been consistently identified as key genetic risk pathways, and risk variants are heavily enriched in or near genes expressed by CD4+ T cells [1,2], particularly in activated cells [3]. Expression quantitative trait (eQTL) studies of activated CD4+ T cells have demonstrated the impact of risk variants on gene expression, with many variants having effects specifically in activated conditions [4,5].

T cells require multiple activation signals from antigen presenting cells (APCs), including a primary signal of antigen recognition (via the T cell receptor, TCR) and secondary signal called costimulation (via costimulatory molecules). Since the 1980s, *in vitro* experimental models of T cell activation have commonly used agonistic antibodies against CD3 to stimulate the T cell receptor and against CD28 (first called Tp44) to provide costimulation [6]. The CD28 ligands, CD80 and CD86, are expressed by dendritic cells (DCs), monocytes and even other T cells in a cell-subset and context-specific way, and changes in their expression impact both the strength of activation and the cell subsets T cells subsequently differentiate into [7-9].

However, beyond CD28, many other molecules can also provide costimulation or coinhibition via other receptor/ligand pairs expressed on T cells and APCs, respectively. The particular set of ligands and receptors that are involved in the interaction are determined by the cell subset, tissue context and cell history of the T cell and APC [10,11]. This can have important implications for disease treatment, as certain T cell subsets evade the CD28-blocker belatacept [12], likely due to the availability of alternative costimulation.

Costimulation pathways were identified as key pathways in the genetics of immune and inflammatory disease early on in GWAS [13]. The list of inflammatory bowel disease risk loci alone [14] includes variants mapped to costimulatory receptors on CD4+ T cells (CD6, CD27, CD226, CD137) and to costimulatory ligands on APCs (CD40, ICOSL). CD28 itself also showed evidence of association for a range of diseases [15]. Some locus-specific research has investigated whether risk variants in alternative costimulatory molecules inhibit their ability to costimulate T cells (e.g. CD6 [16], CD226 [17]), with the most striking example being a 2-fold increase in IL-17 production in CD4+ T-cells from CD226-risk-variant carriers compared to non-risk-variant carriers after CD226 costimulation.

It is known that many T cell genes and pathways are impacted by the level of CD28 costimulation [18], and by the cytokine applied during stimulation [19], and it is likely that differences in other costimulation molecules have similar impacts. Previous work on the transcriptional response to alternative costimulation molecules in human [20] and mouse cells [21] using gene expression microarrays has shown that the same core activation pathways tend to be upregulated by most costimulatory molecules, but that individual costimulatory molecules also often have biases or unique genes or pathways.

It is likely that standard *in vitro* studies of T cell costimulation miss or understudy pathways, genes and gene regulatory elements that are regulated specifically or predominantly by alternative (i.e. non-CD28) costimulatory pathways, and in particular by inflammatory-risk-associated costimulatory genetic variation. For instance, eQTL studies may have lower sensitivity to detect the impact of risk variants on activation-responding genes if they lie in enhancers that are underactive in CD28-dependent costimulation compared to alternative costimulation, or are dependent on regulatory pathways or transcription factors that lie downstream of alternative costimulatory molecules.

In this study, we set out to use modern functional genomics techniques to assess the difference in gene expression, activity of gene regulatory elements and pathway activity between standard CD28-induced costimulation and alternative genetic-risk-implicated costimulatory molecules. We had two main questions: Firstly, how many genes and regulatory elements are preferentially regulated by non-CD28 costimulation, and what sort of functions or pathways are they associated with? Secondly, do we find evidence that genetic risk for inflammatory disease is present and/or enriched in any genes downstream of these costimulatory molecules?

We selected four costimulatory molecules that have previously been identified as candidate causal genes for inflammatory bowel disease, two of which have been well studied in the genome-wide costimulation-induced expression papers reviewed above (CD28 and ICOS), and two not previously studied (CD6 and CD27).

## Results

### Experimental set-up

We used an experimental approach that sorted live CD4+ naive and memory T cells (removing regulatory T cells, Tregs) from N=5 donors, activated them for 24 hours using biotin-conjugated agonistic antibodies bound to streptavidin plates [22], sorted out activated (CD69+ cells) and carried out RNA-seq and ATAC-seq assays to measure gene expression and chromatin accessibility (Figure 1A). The four antibodies were titrated to give comparable levels of activation (measured via CD69 expression, Figure 1B), though in practice CD27 was unable to achieve high levels of activation in memory cells.

**Figure 1:**
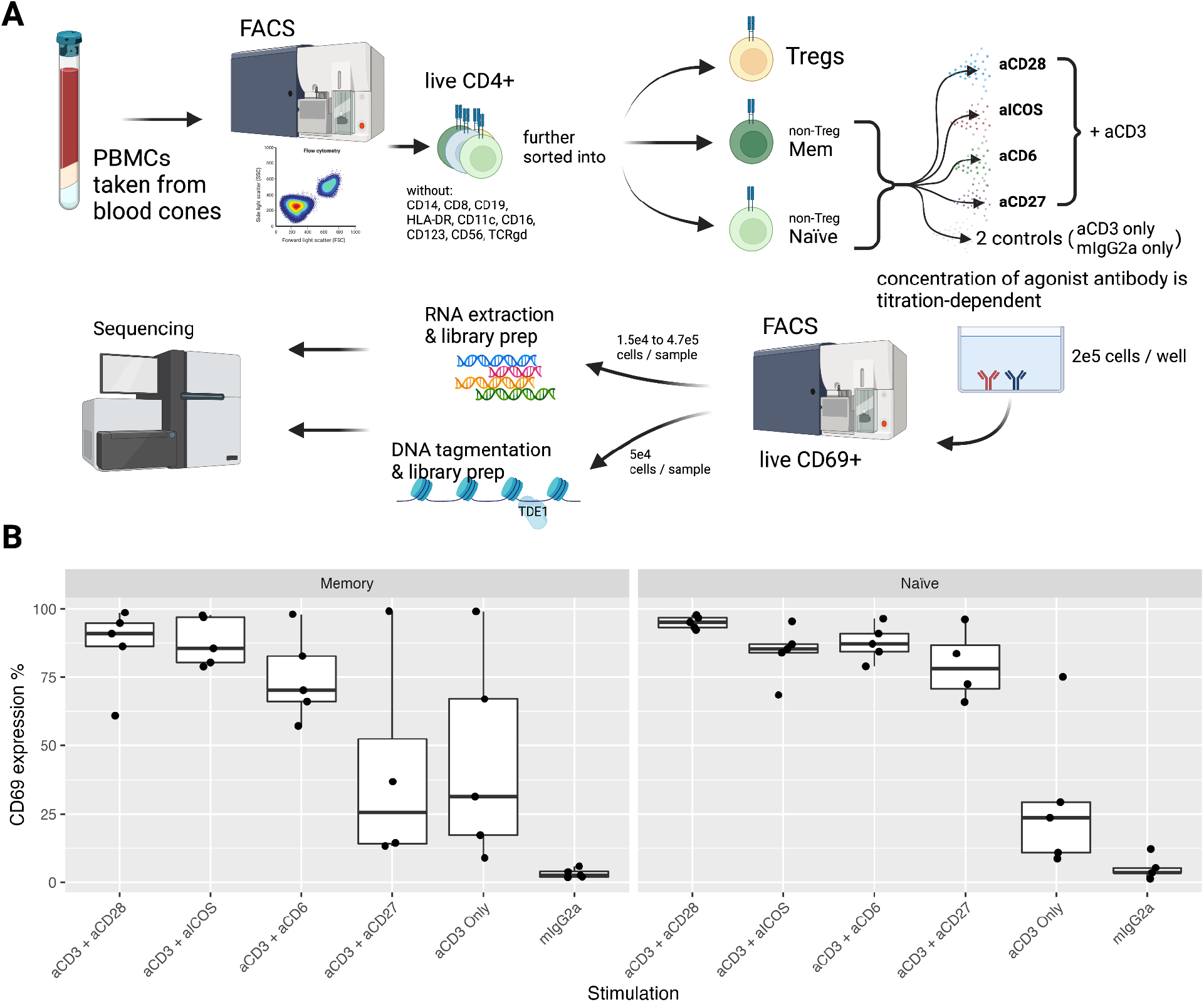
*In vitro* activation and phenotyping of CD4+ T cells under varying costimulation conditions. **A)** Overview of the experiment. **B)** Activation rates of sorted naive and memory CD4+ T cells, measured by expression of CD69 determined by flow cytometry, for different costimulation conditions. mIgG2a = IgG2a isotype control.

For RNA-seq, 58 samples were sequenced (after two failed library QC), with coverage per sample ranging from 22.4M to 50.2M mapped reads. For ATAC-seq, 57 samples were sequenced with coverage per sample ranging from 163.6M to 473.2M mapped reads.

### Overall changes in gene expression and regulation

Principal component analysis demonstrated that the type of costimulation molecule used was the primary driver of genome-wide gene expression across all 58 samples (Figure 2A): the first principal component distinguishes different forms of costimulation, with control samples (cultured with a non-binding IgG2a isotype control antibody) and non-costimulated (cultured with anti-CD3 alone) on one end, standard CD28 costimulation on the other, and CD6, CD27 and ICOS costimulation falling in the middle. The second principal component separated naive from memory cells.

**Figure 2:**
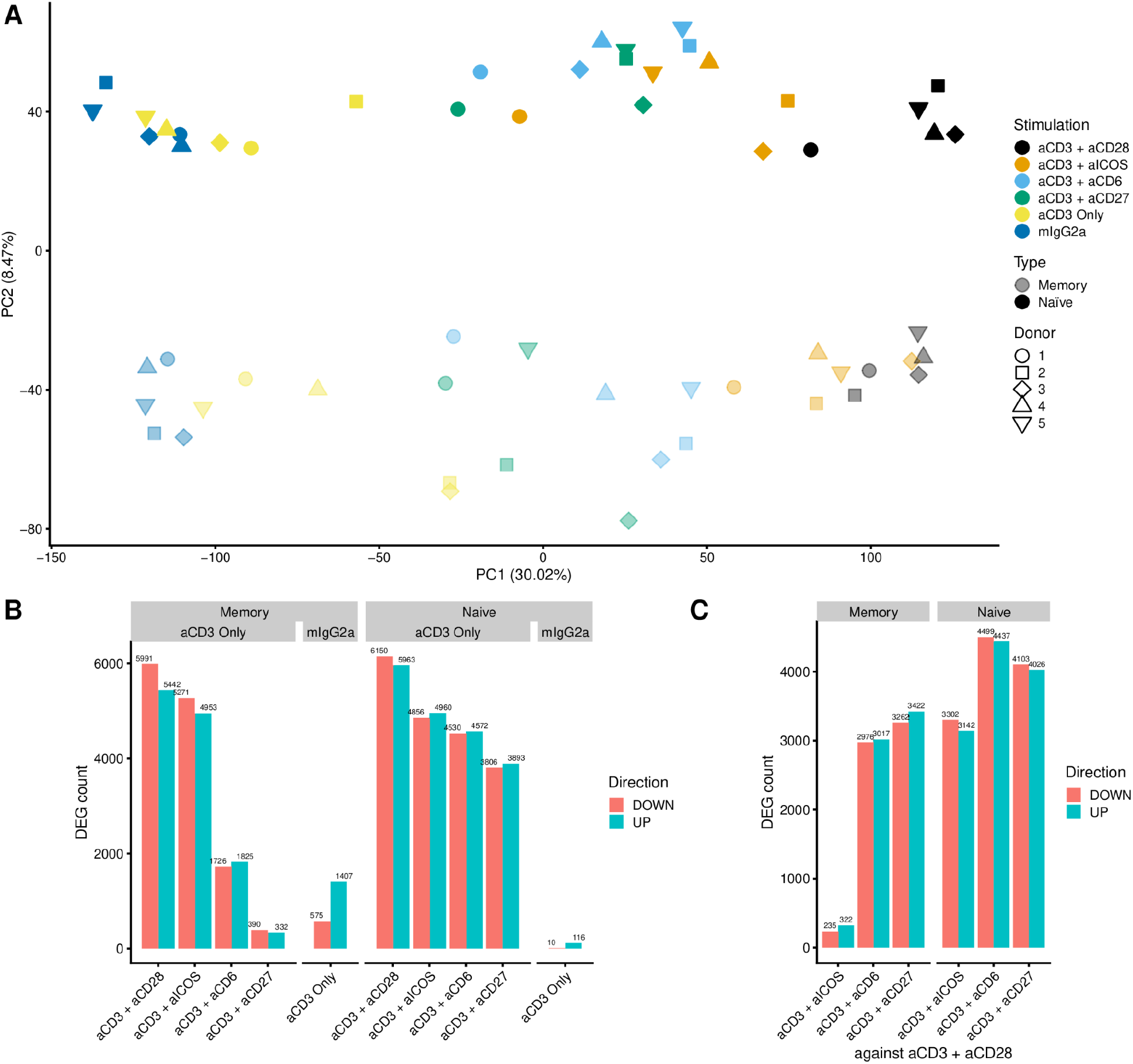
Global changes in gene expression under different costimulation conditions. **A)** Principal component analysis of RNA-seq samples, coloured by stimulation condition and shaded by memory/naive subset. **B)** Count of differentially expressed genes between costimulation conditions and control conditions, broken down by UP- vs DOWN-regulation, naive/memory subset and control condition (shown in gray boxes above the plot, costimulation conditions are compared to aCD3-stimulated-only controls, and aCD3 is compared to mIgG2a isotype controls). **C)** Count of differentially expressed genes between alternative costimulation conditions (aCD6, aCD27 and aICOS) and the aCD28-costimulated condition, broken down by UP- vs DOWN-regulation and naive/memory subset.

We tested for differential expression using DEseq2 [23]. Very large numbers of genes were differentially regulated by costimulation in naive cells, ranging from 6150 upregulated and 5963 downregulated by CD28 to 3806 upregulated and 3893 downregulated by CD27 (Figure 2B). For most genes, the log fold change in gene expression was correlated across the different costimulation molecules (Figure S1), representing the core set of T cell activation genes. The overall strength of the impact of costimulation on gene expression was largest for CD28, and the effect on the majority of genes in the alternative costimulations can be viewed as a “scaled down” version of the impact of that seen in CD28.

To identify genes that show proportionally larger changes in particular alternative costimulatory molecules, we used two different approaches to define “costimulation-biased” genes. The first, which we called the “shape-based method”, selected genes that were significantly upregulated in the alternative costimulation compared to both CD3-alone and CD3+CD28 (or were significantly downregulated in both, Figure S1). The second, which we called the “linear model-based” or “LM-based” approach, tested whether genes had a significantly larger or smaller log fold change than would be predicted based on the log fold change in CD3+CD28 and the genome-wide slope of the regression line (i.e. identifying genes that had significantly larger or smaller effects than would be expected based on a simple change in costimulation strength, Figure S2). The alternative costimulatory molecule with the largest number of costimulation-biased genes was CD6, which had 843 shaped-based and 2279 LM-based genes in the naive condition (Table 1). Overall, there were many more costimulation-biased genes in naive than in memory cells, likely due to the relatively smaller effect of alternative costimulation in memory cells.

**Table 1:** Number of costimulation-biased genes in memory and naive CD4+ T cells under different alternative costimulation conditions compared to CD28 costimulation, estimated using two different methods (LM-based and shape-based).

| Alternative costimulation condition | Memory |  | Naive |  |
| --- | --- | --- | --- | --- |
|  | LM-based | Shape-based | LM-based | Shape-based |
| CD27 | 21 | 50 | 1224 | 492 |
| CD6 | 17 | 69 | 2279 | 843 |
| ICOS | 52 | 77 | 645 | 367 |

In overall genome-wide pattern, gene regulation, measured by accessibility in called peaks, showed a similar pattern to gene expression on its principal components, differentially accessible peaks and costimulation-biased peaks (Supp Figure S3).

### Pathways and transcription factors characterizing alternative costimulation

By applying the fast GSEA [24] method to the LM-based Z scores, we tested for enrichment of Hallmark [25] and KEGG [26] terms among genes relatively upregulated in each alternative costimulation compared to CD28 (Figure 3A and B). A number of effector pathways were downregulated in these alternative costimulations (i.e. upregulated in CD28), including proliferation pathways (G2-M checkpoint genes, E2F targets and cell cycle genes) and, for CD6 and CD27 in naive cells, cytokine response pathways (IFNg and IFNa response genes). Pathways that were characteristic of alternative costimulation include oxidative phosphorylation and lysosome pathways.

**Figure 3:**
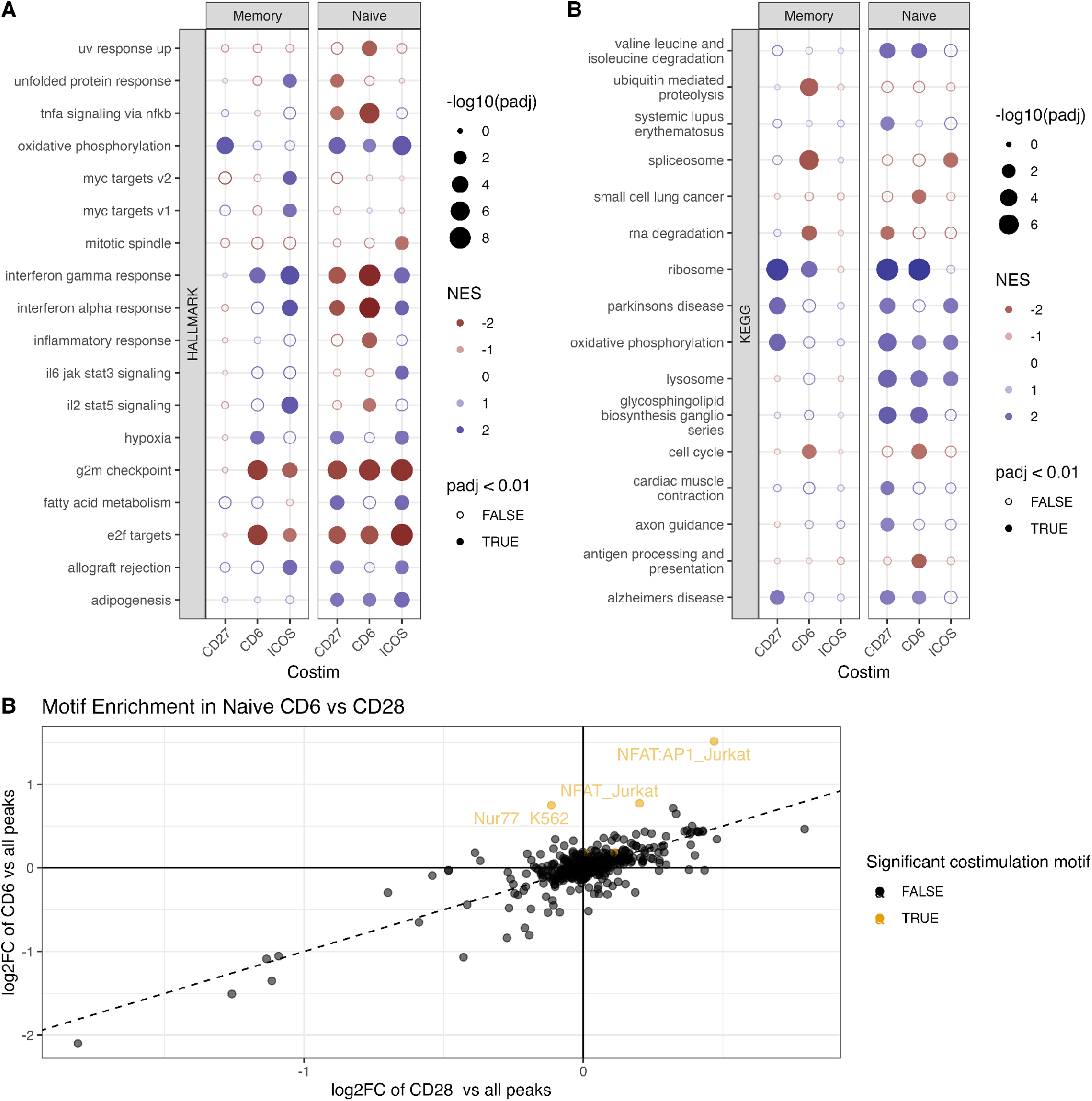
Pathway and transcription factor enrichment analysis. **A)** Hallmark and **B)** KEGG terms significantly enriched (FDR < 0.01) in costimulation-biased genes in at least one alternative costimulation condition NES=Normalized Effect Size, padj=Benjamini-Hochberg adjusted q-value **C)** Enrichment of transcription factor motifs in the CD6 and CD28 costimulation conditions compared to CD3-alone. Yellow dots indicate significant costimulation motifs, defined as motifs which are significantly enriched (FDR < 0.05) both in peaks that are upregulated in the CD6 condition compared to all peaks and compared to peaks that are upregulated in either the CD6 or CD28 conditions.

To investigate the DNA binding proteins driving differences in differential accessible peaks, we tested for enrichment of transcription factor motifs using HOMER [27]. As with gene regulation, most transcription factors were equally active in all costimulation conditions. However, using the same shape-based approach, we found a number of transcription factors differentially active in alternative costimulations, including NFAT, the NFAT:AP1 complex, and Nur77 in CD6 (Figure 3C), the last of which seemed like a unique binding signature in CD6 costimulation and not present in CD28 costimulation. The Nur77 gene (*NR4A1*) was not upregulated in CD6 compared to CD28, suggesting that its activity is not driven by an increased production, but two NFAT genes (*NFATC1* and *NFATC3*) were upregulated in the CD6 condition relative to the CD28 condition.

We carried out further bioinformatic analyses to validate and follow up on the candidate hypotheses suggested by the network analysis. We first tested effector functions (proliferation, cytokine production), which the pathway analysis suggested were lower in alternative costimulations. To validate the impact of costimulation on proliferation, we scored each sample for a cross-species measure of cell proliferation [28] (Figure 4A), demonstrating a clear effect in both memory and naive cells of CD6 and CD27 (and, in naive cells, ICOS) costimulation leading to lower proliferation.

**Figure 4:**
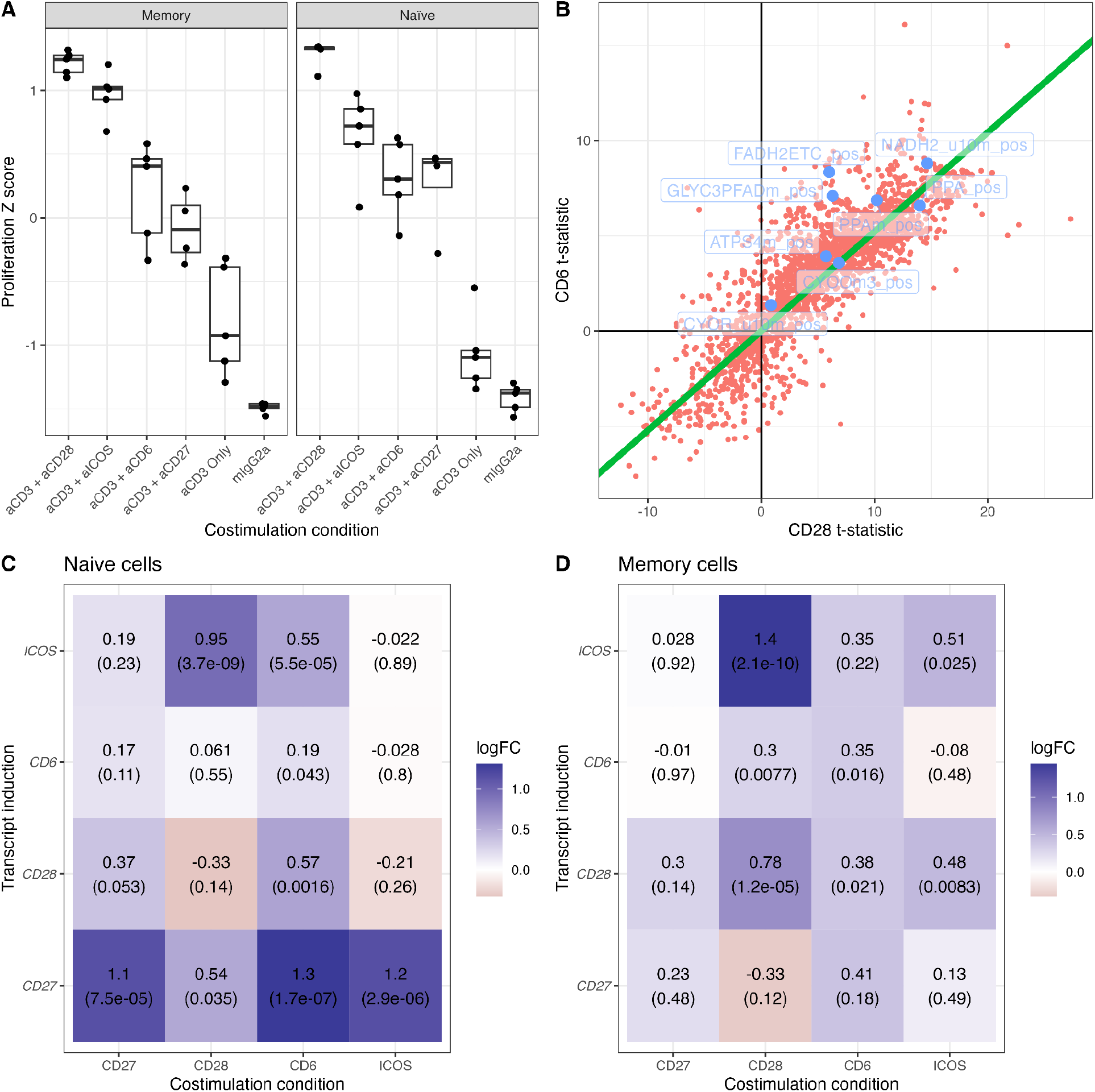
Computational follow-up of key pathways. **A)** Estimated proliferation based on conserved proliferation gene signature for each costimulation condition and cell type. **B)** Differential activity statistics for inferred metabolic rates after costimulation with CD6 or CD28. Green line is the estimated fit, OxPhos reactions highlighted in blue. **C**,**D)** Induction of costimulation molecule transcripts in naive (**C**) and memory (**D**) CD4+ T cells under different costimulation conditions. Text in the boxes are in the form “log fold change (p-value).”

We next turned to the pathways that were enriched in alternative costimulation conditions. To follow up on the lysosome gene set enrichment, we visualized the log fold changes (Figure S4), demonstrating clear signals of upregulation in CD6 across a range of lysosome degradation components, including proteases, glycosidases, sulfatases and phosphatases. The role of the lysosome in costimulation was hard to establish; it was not explained by increased autophagy (LC3 transcripts were not upregulated, and the autophagy KEGG pathway was not enriched), or by antigen presentation (HLA alleles were not upregulated, the antigen processing marker *TAP1* was downregulated in CD6 compared to CD28, and genes in the KEGG antigen presenting pathway were downregulated).

To investigate the role of oxidative phosphorylation metabolism, we used the COMPASS method [29] to infer metabolic reaction rates across 7,440 reactions for each of our samples, and found that, as predicted, OxPhos reactions were more highly upregulated in CD6 compared to CD28 than would be expected under the trend line (Figure 4B). There was no corresponding downregulation in glycolysis reactions in CD6, suggesting that this is not a simple inverse case of oxphos-to-glycolysis metabolic switching as seen in differentiating T cells [30].

For the production of cytokines, we found that almost all cytokine genes were either expressed at a lower level in the alternative costimulation (e.g. *TNF, IFNG* in CD6) or were not expressed at all (e.g. *IL1* and *IL17* in CD6) (Figure S5). The single exception to this rule was the production of XCL1 and XCL2 in naive cells, which was expressed specifically in response to ICOS and no other costimulatory molecules. For cytokine receptors, the pattern was more mixed, with some cytokine receptors expressed at lower levels in alternative costimulations (*TNFR, IFNAR2* in CD6), whereas others expressed at higher levels (e.g. *CCR11*).

We also investigated other genes towards the top of our list. *CD9* stood out as a candidate that was highly expressed by naive cells in response to all costimulation conditions except CD28. While CD9 is involved in the creation of exosomes [31], we did not find other exosome components upregulated in alternative costimulations. Instead, this molecule seems to follow a pattern of cell adhesion genes being upregulated in alternative costimulations, with both the “biological adhesion” and the “leukocyte cell-cell adhesion” GO terms enriched in CD6, CD27 and ICOS, and including a number of adhesion genes (*CD99, LYPD3, CD96*).

We also tested the extent to which costimulatory molecules coregulate each other (Figure 4C). The primary finding was that the *CD27* transcript in naive cells was strongly upregulated by all alternative costimulation molecules (particularly CD6), but weakly, if at all, by CD28. By contrast, the *ICOS* transcript is induced most strongly by CD28, particularly in memory cells, though it is also induced more weakly by CD6. By contrast the *CD6* transcript was only weakly regulated by any costimulation.

### The role of alternative costimulation in genetic risk of inflammatory bowel disease

The ATAC-seq open chromatin data allowed us to test whether regions of the genome that were regulated by different costimulation molecules were enriched for genetic risk of inflammatory diseases. Testing for enrichment of inflammatory bowel disease (IBD) SNP heritability using partitioned LD score regression (LDSC [32]), we found significantly higher heritability in peaks that were upregulated in activated, costimulated naive T-cells compared to CD3-stimulation alone (Figure 5A). This was equally true regardless of the costimulation molecule used. We also tested whether peaks that were shared or unique across both CD28 and alternative costimulation conditions were enriched for IBD risk (Figure 5B), and found the strongest enrichment of risk in peaks that were induced by both, with the second largest enrichment in CD28-specific peaks, and the weakest in alternative costimulation-specific peaks.

**Figure 5:**
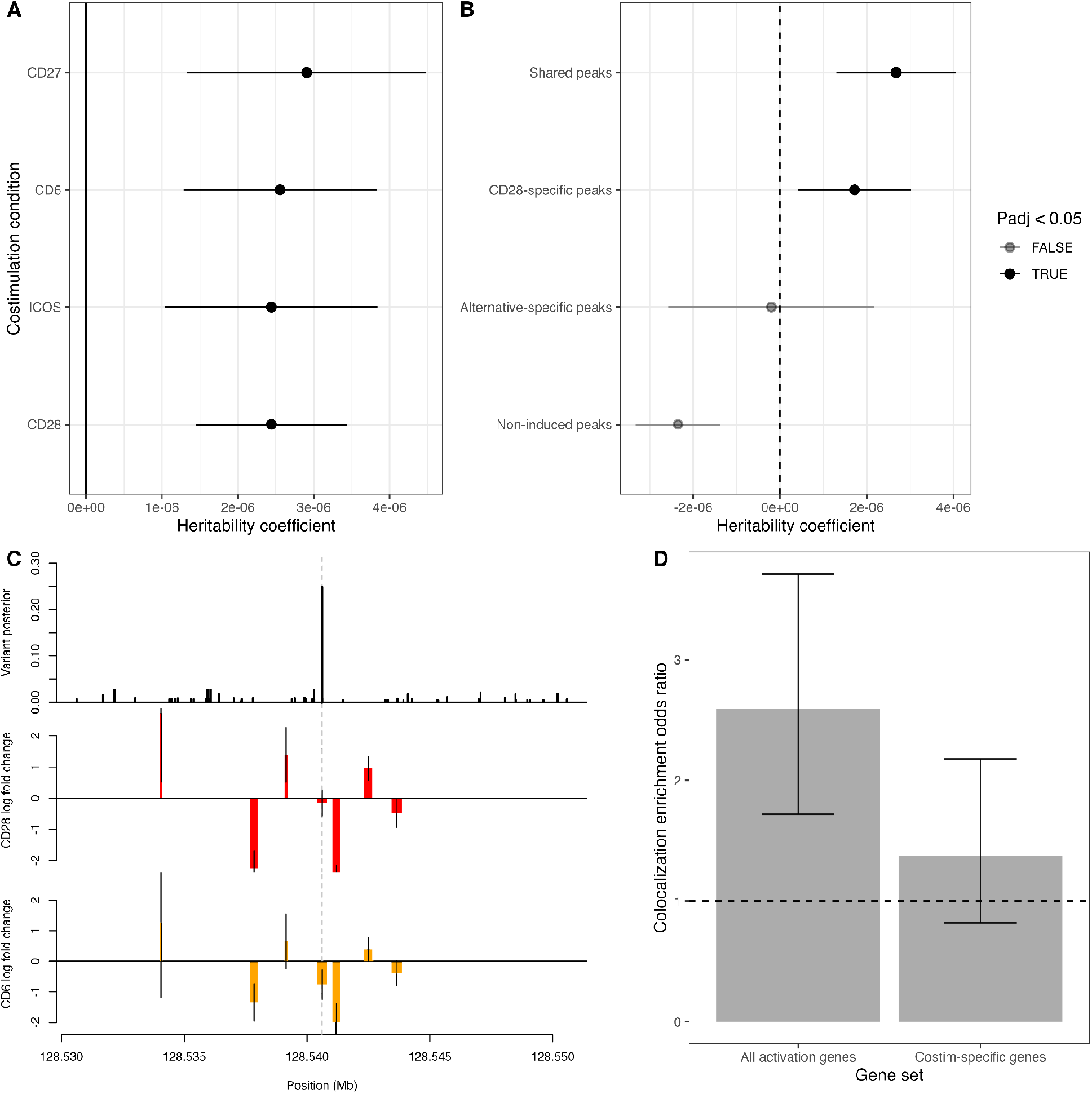
The impact of alternative costimulation on risk variants for inflammatory bowel disease. **A)** Enrichment of heritability in open chromatin regions upregulated in different costimulation conditions (coefficient with 95% CIs). **B)** Enrichment of heritability in regions specifically upregulated in CD28 or in one or more alternative costimulation conditions (CD28-specific and alternative-specific peaks), in both CD28 and at least one alternative costimulation condition (shared peaks) or not upregulated in activated T cells in any condition (neither). **C)** The 8q24 IBD GWAS locus, causality posteriors for candidate causal variants [1], and estimated log-fold changes (with 95% CIs) in the CD28 and CD6 conditions as compared to aCD3-only control. Dashed line shows the location of the highest posterior variant. **D)** Enrichment of IBD associations colocalizing with gene expression in an eQTL study of CD28-costimulated CD4+ T cells [4] among genes upregulated during activation (in any costimulation condition) or specifically upregulated by alternative costimulation and not by CD28.

We intersected our costimulation-biased peaks with genetic fine-mapping results for immune and inflammatory disease associations [1,33] to find candidate risk variants that may act through regulatory elements specific to alternative costimulation. We found nine loci where costimulation-biased peaks contained potential causal variants for inflammatory disease, including seven for inflammatory bowel disease, one for multiple sclerosis and one for primary biliary cirrhosis (Supplementary Table S1). One example of an alternative costimulation-specific risk locus is in the IBD-associated 8q24 gene desert [34], where the lead risk variant is contained within an accessible element that is repressed by CD6 costimulation but not by CD28 costimulation (Figure 5C).

We also intersected high-confidence candidate risk genes for IBD (L2G score > 0.5 [35]) with our costimulation-biased gene list. We looked for genes that are either upregulated specifically in alternative costimulation (i.e. not differentially expressed in the CD28 condition), or had opposite effects in CD28 vs alternative costimulation, as these were genes that would be more likely to be unmapped or give false direction of effect in eQTL studies. Of 76 high-confidence IBD risk genes, 8 met these criteria (Table 2). Of these, five were specific to alternative costimulations, and three went in opposite directions between standard and alternative costimulation. Six of the eight were induced or repressed by CD6.

**Table 2:**
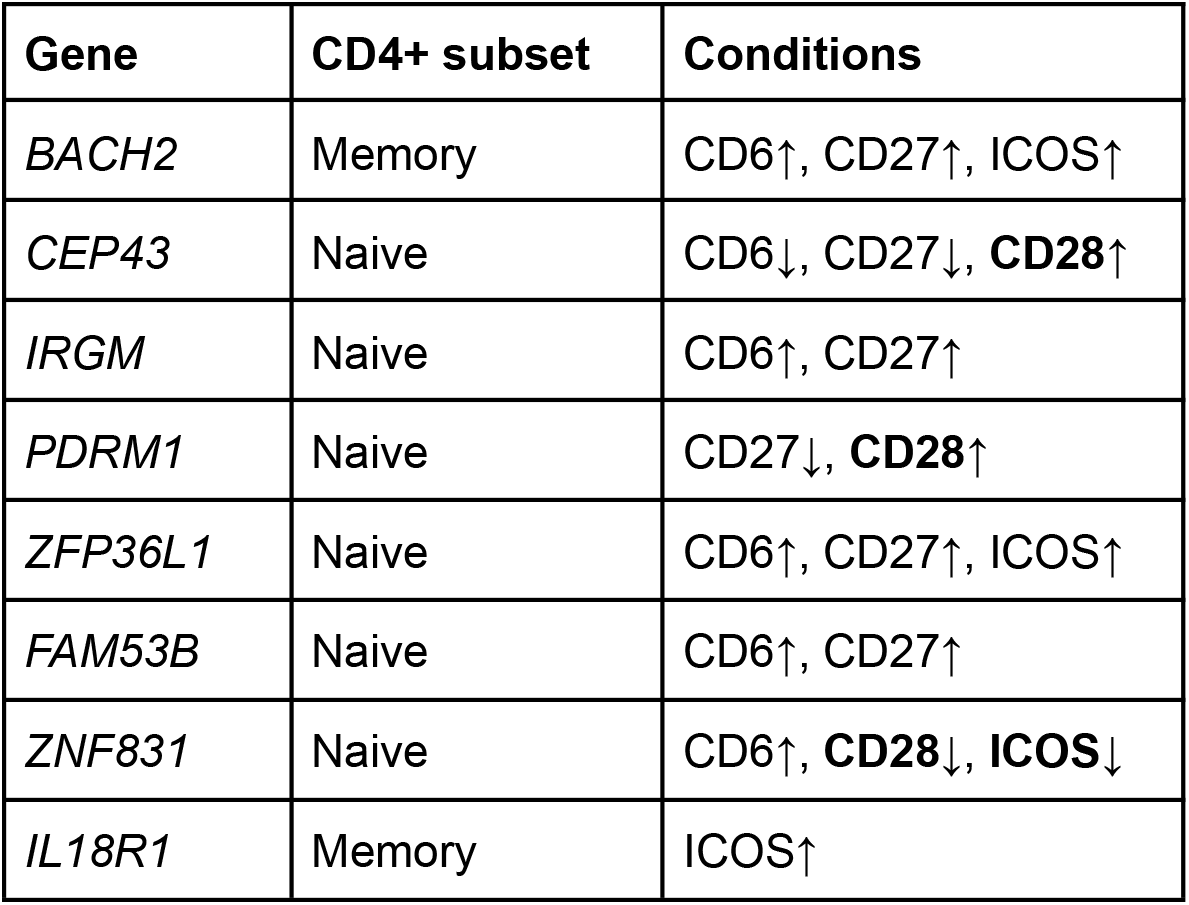
Costimulation-specific inflammatory bowel disease risk genes. High-confidence IBD risk genes that were significantly up- or down-regulated only in alternative costimulation (and not in CD28-costimulated cells), or detected with opposite direction of effect between CD28 and at least one other costimulation condition. ↑/↓ indicate that this gene was significantly upregulated/downregulated in this costimulation compared to the aCD3-alone condition.

We hypothesized that these alternative-costimulation-biased risk genes would be harder to successfully map in traditionally activated (i.e. CD28-costimulated) eQTL studies, and we tested this hypothesis in a recent single-cell eQTL map of CD28+CD3 stimulated CD4+ T cells [4]. Of our 8 candidate IBD risk genes, only one (IL18R1) had a colocalizing eQTL in the CD3+CD28 eQTL dataset (compared to 15/68, or 22%, of the remaining risk genes). Genome-wide, costimulation-specific genes have a lower (and non-significant) enrichment of successfully colocalised IBD risk loci (OR = 1.37, 95% CI = 0.82 - 2.18) than T cell activation genes in general (OR=2.59, 95% CI = 1.72 - 3.71, Figure 5D), providing further evidence that alternative costimulation-specific risk genes are harder to map in standard eQTL studies.

### Experimental validation of key findings at the protein level

We validated a number of our key findings using orthogonal experimental assays of protein expression.

We measured the surface expression of CD6, CD27 and ICOS after CD28 and CD6 costimulation using flow cytometry. We validated that CD27 is upregulated by CD6 compared to CD28 in naive (but not memory) T cells, and that ICOS is more strongly induced by CD28 than CD6 (Figure 6A). As predicted CD6 was not impacted by CD28 costimulation, though it was almost entirely lost from the surface after CD6 costimulation (an effect of ligation-dependent cleavage which has been described previously [36]).

**Figure 6:**
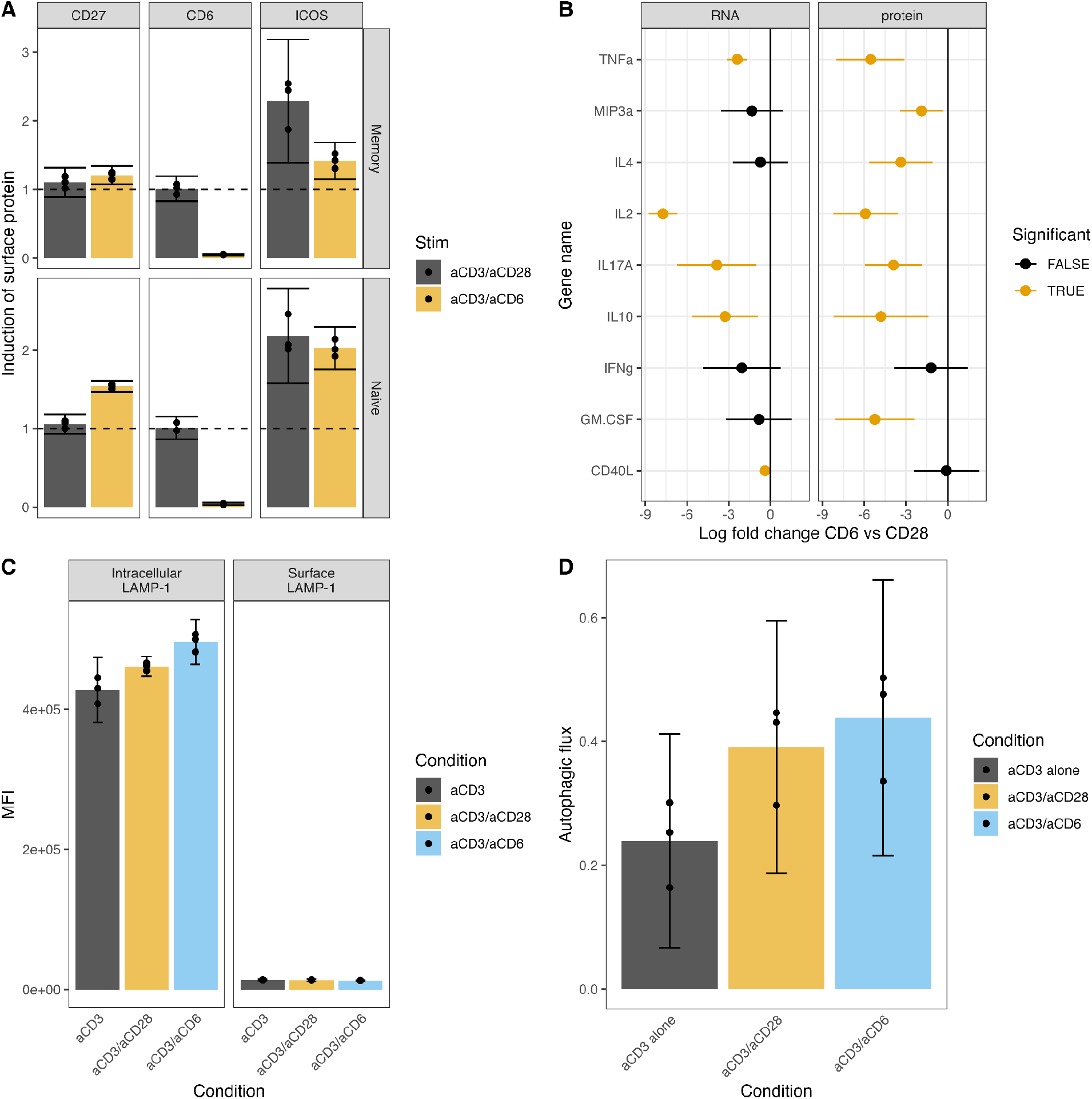
Experimental validation of key findings. **A)** Induction of costimulation receptors on the cell by CD28 and CD6 costimulation, measured by the ratio of mean fluorescence intensity (MFI) in costimulated cells to the MFI in the CD3-alone condition. **B)** Relative change in RNA levels (measured by RNA-seq) and of protein concentration in supernatant (measured by a multiplex immunoassay) between CD6 and CD28 costimulated cells. **C)** Quantification of intracellular lysosome abundance and lysosome secretion under different costimulation conditions, measured using the MFI of intracellular and extracellular LAMP-1. **D)** Quantification of autophagic flux in different costimulation conditions.

We validated the reduced effector function of alternative costimulated cells using a multiplex cytokine assay, showing either absence or significantly reduced secretion of key effector cytokines by CD4+ T cells after costimulation with CD6 compared to CD28. The level of reduction in protein concentration was consistent with the level of reduction of gene expression (Figure 6B).

Finally, we validated the increase in lysosomes using intracellular staining of LAMP1 measured by flow cytometry, demonstrating that T cells accumulate lysosomes after CD6 costimulation compared to CD28 costimulation (Figure 6C). We further investigated whether this increased lysosome formation could be explained by increases in either autophagy [37,38] or lysosome exocytosis [39], but saw neither evidence of lysosome secretion or an increase in autophagic flux after 24 hours of CD6 costimulation (Figure 6C,D).

## Discussion

We have carried out a detailed assessment of the different effects genetic-risk-associated costimulatory molecules play on gene regulation, gene expression and downstream pathways in activated CD4+ T cells. This has revealed a number of key findings about the functional impacts of these signaling molecules, and the role they may play in the genetics of inflammatory disease. We have also successfully validated a set of these findings at the protein level using orthogonal experimental approaches.

The majority of the changes downstream of T cell activation, as has been noted by previous research, are activated (to some degree) regardless of the specific costimulation molecule used, though usually with a larger effect in the CD28 condition. Costimulation with CD28 has a more dramatic impact on T cell function than other costimulators across three levels of the signaling cascade. Firstly, anti-CD28 is a more potent costimulator than other molecules, and in our experiments we required 6.7x the concentration of anti-CD6 to achieve a comparable level of T cell activation (measured by CD69 positivity). Next, even at comparable levels of activation, and looking at only activated (CD69+) cells, we saw a larger degree of chromatin remodeling and a more dramatic impact on gene expression in the CD28 condition than for other costimulation molecules. Finally, even after controlling for the genome-wide patterns, we found that genes underlying effector functions, such as proliferation and cytokine release, were further enriched in CD28 compared to other costimulators, relative to other genes.

There were numerous genomic regions, genes and pathways that diverged from this overall trend and responded exclusively or predominantly to alternative costimulation. This was particularly true in naive cells, and particularly during CD6 costimulation. Alternative costimulation called T cells to have a relative shift towards what appears to be a “getting ready” cell state. These cells secrete less cytokine and proliferate less. They begin to overexpress genes that may play roles in searching out further activation signals, including further costimulation (by upregulating *CD27*), homing to inflammatory areas (via *CCR11* [40]) upregulating certain cytokine receptors (*TNFR, IFNAR2*) and increasing activity of activation-mediating transcription factors (Nur77/*NR4A1* [41]). They also undergo a number of intracellular changes, including the overproduction of lysosomes and a relative increase in oxidative phosphorylation. The lysosome overproduction did not appear to be related to autophagy or antigen presentation, and we did not see evidence of secretory exocytosis (though the 24-hour time point may have been too early to detect it). Lysosome-mediated protein degradation plays a role in many other T cell functions [42], some general (e.g. increasing protein turn-over) and some specific (including sensitizing cells to activation by degrading CTLA-4 [43]), and the same is true for metabolic state [44]. Further investigation into these pathways, and how they are altered by genetic risk variants for inflammatory disease, will shed light into the downstream functions of these pathways and their role in immunopathology.

In this study, we were specifically interested in whether previous studies of CD3+CD28 activation of T cells could have missed important genetic risk pathways, for example by failing to find eQTLs in genes expressed only downstream of genetically implicated alternative costimulation molecules. Our heritability analysis suggests that the majority of risk variants lie in general T cell activation genes, and that the alternative costimulation molecules are no better at inducing the expression of risk genes than CD28. In fact, regulatory regions that were only activated by alternative costimulation and not by CD28 had significantly less enrichment of heritability than regions regulated by all costimulatory molecules. However, this does not mean that alternative-costimulation-specific pathways play no role in disease risk: we found both risk variants and risk genes that are exclusively regulated by alternative costimulation, both for inflammatory bowel disease (including *IRGM* and *ZFP36L1* in CD6 costimulation) and for psoriasis (*XCL1/2* in ICOS costimulation [45]). As predicted, alternative costimulation risk genes are less likely to colocalize with gene expression in CD28-costimulated CD4+ T cells in a recent single-cell eQTL study [4].

There are many limitations of this study, and much more could be done. We picked a 24 hour time-point in order to capture an early pre-proliferative, pre-differentiation state. However, both earlier time-points (to capture the immediate effects of costimulation) and later time points (to capture the effects of differentiation and T cell polarization) could also include valuable information. There are further experimental studies that could and should be done to validate more findings at the protein level, and measure their impacts on downstream cellular functions. There is also further investigation that can be done on the impact of combinations of costimulation - here we have modeled the effect of “pure” alternative costimulation (as might be seen from a CD80/86-negative APC, or a CD28-negative T cell), but in practice a T cell will often receive a combination of costimulation signals, which may have non-additive impacts. We have also used bulk RNA-seq and ATAC-seq, sorted only on activation state and naive vs memory subset, which may miss strong impacts of certain costimulatory molecules on certain rare T cell subsets.

A final limitation of the study is that, while we have adjusted for stimulation strength as much as possible (by achieving comparable levels of CD69-positivity, sorting CD69+ cells and adjusting for global changes in gene regulation/expression), it remains difficult to distinguish between the impacts on T cell function due to a weaker stimulation compared to specific signaling pathways downstream of a given costimulation molecule. The “getting ready” T cell states we describe, induced by alternative costimulation, may in fact be “weaker activation” states, where T cells receive a signal sufficient to activate but not sufficient to commit to full effector status. Ongoing studies of the impact of TCR stimulation affinity on T cell activation will test this hypothesis.

A key hypothesis that we began this study with was that genetic risk variants in alternative costimulatory molecules could be increasing risk by impacting genetic risk pathways that were specifically downstream of these molecules. This is still an active hypothesis, and we have found candidate genes and variants where this may be the case. However, the overall similarity in the downstream effects of CD28 and other risk-associated costimulation molecules also leaves open the possibility that these variants are acting as overall modulators of the shared T cell activation response (with any costimulation-specific effects being coincidental to disease risk). Ultimately, it will require costimulation-specific eQTL maps of CD4+ T cells to answer the question of how costimulation and disease risk interact, by mapping cis-eQTLs for risk variants and genes that are only regulated by alternative costimulation, and by mapping the downstream effects of costimulatory risk variants by finding their trans-eQTLs effects on downstream activation genes.

## Acknowledgements

The authors would like to thank Matthias Friedrich, Claire Pearson, Fiona Powrie, Ghada Alsaleh, Katja Simon, Irina Uldalova, Jelena Bezbradica Mirkovic, Holm Uhlig, Pablo Céspedes, Omer Dushek and Gosia Trynka for providing advice and guidance during this project. AV, JH, MD and LJD were funded by the Kennedy Trust for Rheumatology Research, LJD, JH, CM and MD were funded by the Wellcome Trust (grants 208750/Z/17/Z and 100262Z/12/Z) and KC was funded by the Rhodes Trust. Visual representation of experiment (Figure 1A) created with BioRender.com under license OO24OFTPGD.

## Data and code availability

The gene count matrix for RNA-seq and peak count for ATAC-seq, as well as sample meta-data, differential expression and accessibility results and costimulation-bias test results for all genes and called peaks, are publicly available at this address: https://doi.org/10.6084/m9.figshare.21614028 The code necessary to identify costimulation-biased genes (from the Methods section) is available publicly at this address: https://doi.org/10.6084/m9.figshare.21893466

## Methods

### Experimental methods for generating RNA-seq and ATAC-Seq data

#### Protein biotinylation

Biotinylated antibodies for CD3, CD28 and ICOS were sourced from companies (clone IDs mentioned in the T-cell stimulation section of methods). Human Anti-CD27 agonist antibody (100111-1, AMSBio) and Mouse Anti-CD6 agonist antibody (Clone UMCD6, Sigma Aldrich) were biotinylated using the EZ-Link™ Sulfo-NHS-LC-Biotin kit (ThermoFisher) as per the manufacturer’s instructions. Biotinylated antibodies were purified using Zeba™ Spin Desalting Columns (7K MWCO, 0.5 mL, ThermoFisher). Recovered protein was quantified using Micro BCA™ Protein Assay Kit (ThermoFisher) and biotin levels were quantified using Pierce™ Fluorescence Biotin Quantitation Kit (ThermoFisher) as per manufacturer’s instructions. Purified biotinylated antibodies were stored at -80°C until use.

#### Sample collection and initial T-cell enrichment

Human PBMCs were isolated from blood leukocyte cones obtained from apheresis donations of patients giving informed consent, supplied by the John Radcliffe NHS Blood and Transplant service (ethical approval REC 11/H0711/7). Briefly, PBMCs were isolated using density centrifugation over Percoll. Pelleted cells were then treated with ACK lysis buffer and washed. CD4^+^ T-cells were then isolated using the human CD4^+^ T-cell MojoSort negative magnetic selection kit (Biolegend). Enriched CD4^+^ T-cells were then rested overnight in a 37°C 5% CO_2_ incubator in complete RPMI (RPMI1640, Pen/Strep, 10% FCS, Sodium Pyruvate, HEPES, Gentamicin, Glutamax).

#### Isolating Naïve and Memory T Cells

After overnight culture, naïve and memory T cells were isolated using fluorescence-activated cell sorting using a FACSAria III cell sorter. The cells were gated such that single live (LiveDead Supplier) cells were gated, subsequently CD4^+^ (PE-Cy7, Biolegend) [CD14 CD8 CD19 HLA-DR CD11c CD16 CD123 CD56 TCRgd]^−^ (FITC, Biolegend) cells were gated, Tregs were then isolated on the basis of CD127^lo^ (BV421, Biolegend) CD25^+^ (APC, Biolegend), non-sorted cells were then further sorted into Naïve (CD45RA^+^ (BV785, Biolegend) CCR7^+^ (PE, Biolegend)) and Memory Cells (all cells bar CD45RA^+^ CCR7^+^). Sorted cells were washed and resuspended in freezing media (90% FCS + 10% DMSO) and aliquots were frozen overnight at -80°C then transferred to storage in liquid nitrogen.

#### T-cell stimulation

Naive and memory T-cells were defrosted and washed before stimulation. Activation plates were created by immobilising commercially biotinylated Anti-CD3 (Clone OKT3, Miltenyi Biotec) at 0.1ug/ul with either commercially biotinylated Anti-CD28 (Clone 15E8, Miltenyi Biotec), Anti-ICOS (Clone ISA-3, ThermoFisher), in-house biotinylated Anti-CD6 or in-house biotinylated Anti-CD27 into streptavidin-coated plates (ThermoFisher) at concentrations of 0.1, 0.5, 0.67 and 0.49ug/ul respectively. Control wells were created in the same manner by immobilising commercially biotinylated Mouse IgG2a (Biolegend) alone at a concentration of 0.5ug/ul. Antibodies were incubated for 45 minutes at room temperature on a plate shaker at 1000 rpm. Plates were washed once with sterile PBS. After washing, Naïve or Memory T-cells were added to each stimulation condition (2 × 105 per well in 200μl of cRPMI) in multiple instances. Plates were incubated at 37°C with 5% CO_2_ for 24hrs.

#### Sorting T-cells post-activation

After stimulation, Naïve or Memory cells from the same donor under the same activation condition were pooled and stained for viability (eFluor™ 780 fixable stain) and Fc Blocking in ice cold PBS for 15 minutes. After this time, CD69 (PE-Dazzle 594, Biolegend) in ice cold PBS + BSA + EDTA was added and staining continued for an additional 30 minutes. Cells were then washed and resuspended prior to sorting using a FACSAria III cell sorter. Anti-CD3 only stimulated cells were sorted on the basis of live single cells, control samples (Mouse IgG2a) were sorted on the basis of live single cells that are CD69^−^ (PE-Dazzle 594, Biolegend) and the stimulated samples (anti-CD3 plus either anti-CD28, anti-ICOS, anti-CD6 or anti-CD27) were sorted on the basis of live single cells that are CD69^+^ (PE-Dazzle 594, Biolegend). Sorted post-activation naïve and memory T cells were counted.

#### RNA-seq

Between 1.5 × 10^4^ – 4.7 × 10^5^ cells per sample were isolated and placed into 300μl of Buffer RLT and stored at -80°C until processing for RNA isolation. Samples were thawed at room temperature and RNA extracted using the RNeasy Micro Kit (Qiagen) as per manufacturer’s instructions. Purified RNA was quantified via Nanodrop. The purified RNA was sent to a sequencing center.

#### ATAC-seq

ATAC-Seq was carried out as per a previously published protocol [M1]. 5 × 10^4^ cells per sample were isolated and washed in ice cold PBS then subsequently incubated in cold ATAC-Resuspension Lysis Buffer and triturated then incubated for 3 minutes. The suspension was then washed in plain ATAC-Resuspension Buffer and spun down, the supernatant was carefully removed to leave the isolated nuclei. 50μl of transposition buffer, containing the TDE1 Tagment DNA Enzyme (Illumina), was added to each sample and incubated for 30 minutes at 37°C shaking at 1000 rpm in an Eppendorf shaker. The tagmented DNA was isolated using the MinElute PCR purification kit (Qiagen) as per manufacturer’s instructions. Post-purification, each DNA sample was individually indexed using the Nextera XT Index Kit v2. In total 57 libraries were indexed and pooled. The pooled library was sequenced.

### Experimental methods for follow-up experiments

#### Human CD4+ T cell isolation and culture for follow-up experiments

Human blood leukocyte cones obtained from apheresis donations were supplied by the John Radcliffe NHS Blood and Transplant service. PBMCs were obtained by density centrifugation with Histopaque-1077 (Merke). Pelleted cells were then treated with ACK lysis buffer (Sigma) and washed. Human CD4+ T cells were negatively selected from PBMCs using a Human CD4+ T cell isolation kit (Miltenyi). The purity of CD4+ T cells was assessed using anti-CD3 APC (Biolegend) and anti-CD4 Pacific Blue (Biolegend) by flow cytometry (Aurora, Cytek). After isolation, cells were resuspended at 1 × 10^7 in 90% FBS (ThermoFisher) 10% DMSO (Sigma Aldrich), stored at -80 overnight before long term storage in liquid nitrogen. After thawing, CD4+ T cells were resuspended in complete RPMI (RPMI-1640 (Sigma-Aldrich), 10% FBS (supplier), 1mM sodium pyruvate (ThermoFisher), 2mM glutaMAX (ThermoFisher), 10mM HEPES, 100U/mL Penicillin/Streptomycin) at 1 × 10^6 cells/mL. These cells were cultured for 24 hours with either anti-CD3 biotin (Miltenyi)/anti-CD28 biotin (Miltenyi) or anti-CD3 biotin (Miltenyi)/aCD6 biotin (LSBio) on streptavidin coated 96 well plates (ThermoFisher) to yield 50% CD69 expression (antibody amount required determined by prior experimentation). After 24 hours cells were harvested for downstream analysis and culture supernatant was stored at -80 for later proteomic assays.

#### Surface staining for flow cytometry

Flow cytometry analysis was performed on CD4+ T cells immediately after culture. Viable CD4+ T cells were identified for analysis using eBioscience Fixable Viability Dye eFluor™ 780 (ThermoFisher), anti-CD3 APC (Biolegend), anti-CD4 Pacific Blue (Biolegend). Live CD4+ T cells were analysed using anti-CD69 FITC (Biolegend), anti-CD25 PE (Biolegend), anti-CD45RA Brilliant Violet™ 605 (Biolegend), anti-CCR7 Brilliant Violet™ 711 (Biolegend), anti-CD6 Brilliant Violet™ 510 (BD Bioscience), anti-ICOS PE-Cy7 (Biolegend), anti-CD27 PerCP Cy5.5 (Biolegend). Cells were then fixed with 2% PFA. Cells were acquired (Aurora, Cytek) and analysis was performed using FlowJo version 10.7.1 (TreeStar, USA) and gates set using fluorescence minus one (FMO) controls.

#### Autophagic flux and lysosome assay

PBMCs were obtained and cryopreserved at 5 × 10^7^ as described above. Thawed PBMCs were resuspended at 1 × 10^6 in complete media. PBMCs were cultured for 24 hours with either anti-CD3 biotin (Miltenyi)/anti-CD28 biotin (Miltenyi) or anti-CD3 biotin (Miltenyi)/aCD6 biotin (LSBio) on streptavidin coated 96 well plates (ThermoFisher) to yield 50% CD69 expression on CD4+ T cells (antibody amount required determined by prior experimentation). Autophagy levels were measured using an adapted protocol for the Guava Autophagy LC3 Antibody-based Assay Kit (Luminex). Briefly, autophagy was measured after 2 hour treatment with either bafilomycin A1 (Cayman chemical) or vehicle. Cells were then stained with eBioscience Fixable Viability Dye eFluor™ 780 (ThermoFisher), anti-CD3 Brilliant Ultra Violet™ 395 (BD Bioscience), anti-CD4 Brilliant Ultra Violet™ 805 (BD Bioscience), LAMP-1 APC (Biolegend) and anti-CD69 Brilliant Violet™ 650 (Biolegend). Cells were then washed and permeabilised with 0.05% saponin. Cells were stained with anti-LC3-II FITC (Luminex) and LAMP-1 PE (Biolegend). Cells were then fixed with 2% PFA before acquisition (Aurora, Cytek) and analysis was performed using FlowJo version 10.7.1 (TreeStar, USA) with gates being set using FMO controls.

#### Luminex proteomic assay

The concentration of 17 analytes (CD40 Ligand, GM-CSF, IFN-γ, IL-1β, IL-2, IL-4, IL-5, IL-6, IL-10, IL-12p70, IL-13, IL-15, IL-17A, IL-17E, IL-33, MIP-3α AND TNF-α) were assessed from the culture supernatant of the activated CD4+ T cells by Human Th9/Th17/Th22 Luminex Performance Assay 17-plex Fixed Panel (R&D systems, USA). The assay was performed according to manufacturers instructions and samples diluted 1:2 with calibrator diluent. The plate was read using the Luminex Magpix analyser (R&D systems, USA).

### Computational methods

#### Sequencing data quality checking, alignment and count processing

For RNA-seq data, FastQC v.0.11.9 (bioinformatics.babraham.ac.uk/projects/fastqc/) was used before alignment to check the quality of the reads, with reports over multiple samples compiled by MultiQC, v.1.11 [M2]. Hisat2, v.2.1.0 [M3], was used to align the paired-end reads to the GRCh38 genome. SAMtools, v1.9 [M4] was used to sort and index alignment data. Picard, v.2.10.9 (broadinstitute.github.io/picard/) was used to check for duplicates and a number of other alignment metrics. Aligned reads (BAM output) were used as input for the featureCounts function of the Rsubread library, v.2.0.1 [M5] to obtain the counts matrix.

For ATAC-seq data, pre-alignment QC was performed using FastQC v.0.11.9, and QC reports were compiled using MultiQC v.1.7. Reads were mapped to the GRCh38 reference genome using BWA-MEM v0.7.15 using default parameters. Read duplicates were removed with the “MarkDuplicates” function of Picard v.2.23.0 using “REMOVE_DUPLICATES=true”. Only reads with a length less than 100 bp, with a mapping quality >= 10, and mapping to chromosome 1-22, X, and Y were kept by filtering with the “view” function of SAMtools v.1.12. Reads from shallow and deep sequencing were merged using the “merge” function of SAMtools v.1.12. Mapped reads were QC’ed using SAMtools (“stats”, “flagstat”, and “idxstats” functions) and Picard (CollectInsertSizeMetrics and “CollectAlignmentSummaryMetrics” functions). Peaks were called with MACS2 v.2.2.6 using default parameters and “-f BAMPE -g hg38 --keep-dup all”. For the subsequent processing steps, Bioconductor v.3.14 was used. A non-overlapping set of consensus peaks was found using the “reduce” function of the GenomicAlignments package over all samples. Consensus peaks overlapping with blacklist regions (hg38-blacklist.v2.bed from https://github.com/Boyle-Lab/Blacklist/tree/master/lists [M6]) or present in fewer than 2 samples were removed. Overlaps of mapped reads from all samples with consensus peaks were counted using the “summarizeOverlaps” function with “singleEnd = FALSE” from the GenomicAlignments package.

#### PCA, differential expression and gene ontology enrichment analysis

For RNA-seq data, in order to perform principal component analysis (PCA), the count matrix was normalised to counts-per-million (CPM) and then genes with zero variance were discarded. The resulting matrix was used as input for the *prcomp* function from R, v.4.1.0. Plotting of the results was achieved by adding shape, colour and alpha descriptions (of donor, stimulation and cell-type) to the PC1 and PC2 coordinates with ggplot2, v.3.3.5. For analyses of differential expression, the counts matrix was processed with DESeq2, v.1.32.0 [23], separately for memory and naive samples, with the alpha threshold of 0.05, with all other parameters set to default and the design matrix set as ∼ *stimulation*.

For ATAC-seq data, PCA analysis was performed using the “plotPCA” function after a variance stabilizing transformation was applied to count data using the “vst” function, both from the DESeq2 package, v.1.34.0. Differential accessibility analyses were performed with DESeq2 using default parameters, Benjamini-Hochberg p-value adjustment with an alpha of 0.05, and a design formula of “∼stimulation.” Peaks were annotated using the “annotatePeak” function from the ChIPseeker package, v.1.30.3, with “TxDb = TxDb.Hsapiens.UCSC.hg38.knownGene, annoDb=org.Hs.eg.db.”

#### Identifying rebels against the stimulation-strength trend

##### LM method

One method of removing the effect of stimulation strength from other costim effects leverages regression. A univariate linear model is fit that predicts the log of the fold-change (logFC) of non-canonical costimulation (when compared to CD3-only) from the logFC of canonical costimulation (also when compared to CD3-only). The residuals of this univariate model indicate how much an RNA deviates from the typical stimulation strength-dependent effects, if the standard error of the linear model is taken into account:

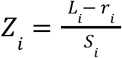

where ***i*** indexes all the RNA transcripts, ***L***_***i***_ represents the log fold-change of this RNA between a non-canonical costim and CD3-only, ***r***_***i***_ represents this RNA’s residual in the linear model predicting ***L*** from correspondent log fold-changes between CD28 and CD3-only, and ***S***_***i***_ represents the following equation:

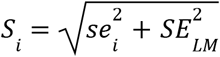

where ***i*** indexes all the RNA transcripts, ***se***_***i***_ is the standard error of the ***L***_***i***_ and ***SE***_***LM***_ is the standard error of the linear fit. Two-sided P-values are obtained from said Z-scores and then are adjusted for multiple testing with Benjamini-Hochberg.

##### “Shape” method

To select a gene as an non-canonical costim-specific gene (for example, CD6-specific), the “shape” method is based on two rules, either the gene is:

1. Significantly upregulated in non-canonical v. CD3-only **and** significantly upregulated in non-canonical v. CD28.
2. Significantly downregulated in non-canonical v. CD3-only **and** significantly downregulated in non-canonical v. CD28.

An intuitive sketching of the methods can be seen below:

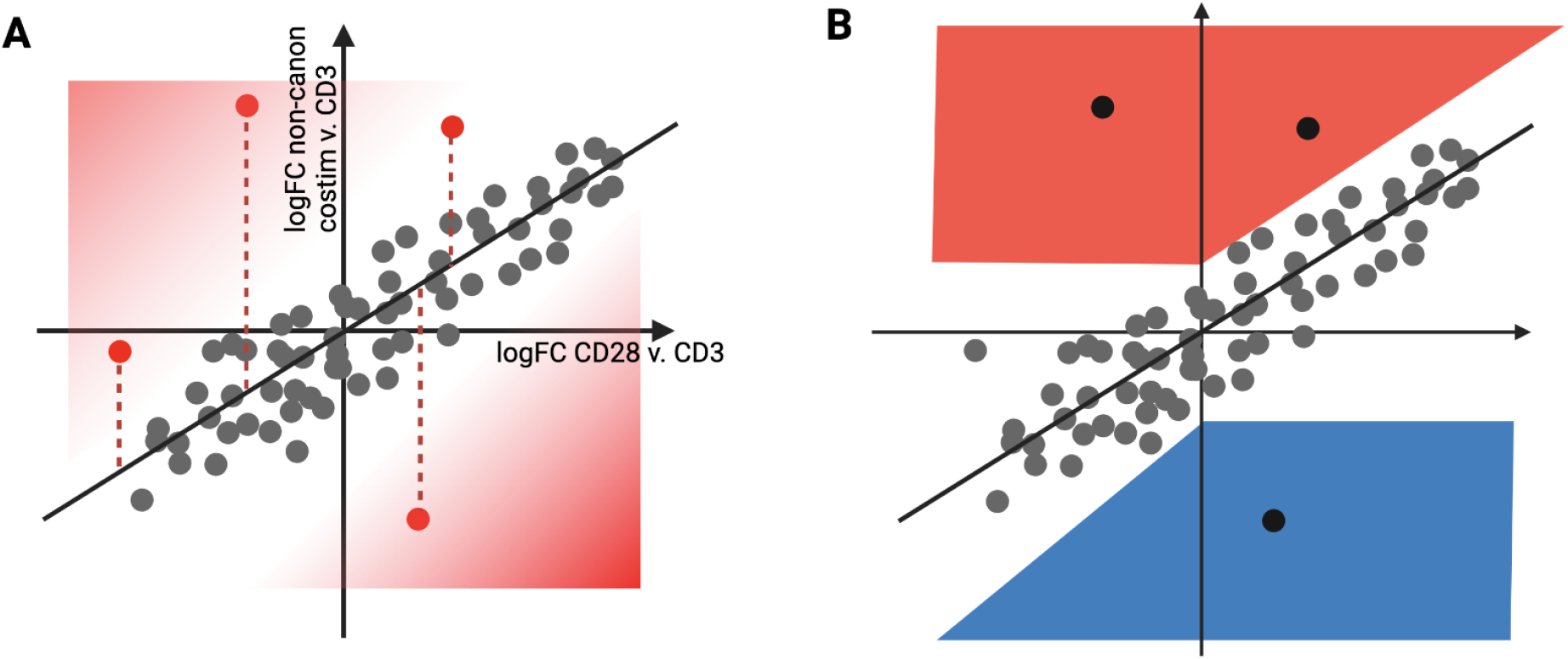

***Example stimulation-strength trend and the two methods of selecting genes that rebel against the trend. A)*** *The linear method of identifying costim-specific genes involves regressing the logFCs of one costim on the other and then extracting significant Z scores based on the residuals of each transcript and the model’s standard error*. ***B)*** *The “shape” (rule-based) method involves a set of rules to select the genes that are costim-specific based on just the output of DESeq2. In a certain way, it is more optimal than the model in figure A because it does not select the leftmost outlier selected by the LM*.

#### Pathway enrichment tests

We measured enrichment using the R package FGSEA, version 1.18.0 [24], with gene-set information extracted from the msigdbr R package, version 7.4.1 for GO, Hallmark and KEGG terms. The ranks supplied for FGSEA trend-bucker analyses were the Z scores of the transcripts, obtained from their residuals in the *logFC*_*non-canon costim*_ *∼ logFC*_*CD28*_ model. KEGG plots were made with pathview, v1.32.0 [M7].

To estimate proliferation rates from gene expression data, we used a previously described method [28] that tests for enrichment of 370 conserved proliferation genes using single sample GSEA (ssGSEA [M8]). ssGSEA to estimate proliferation rates, which is used in this analysis as well. We used the publicly available implementation of ssGSEA on Rpubs by Pranali S. (https://rpubs.com/pranali018/SSGSEA).

#### Metabolic state estimation using COMPASS

To evaluate OxPhos changes, we used COMPASS (version 0.9.10.2), a software designed to estimate metabolic parameters in single-cell and bulk RNA-seq datasets [29]. Raw reaction consistency values output by Compass were then processed as a negative log and close-to-constant reactions were then dropped. All parameters left on default settings.

#### GWAS enrichment testing

Enrichment testing of european GWAS summary statistics of IBD [14] was carried out using LDSC [32], using parameters and implementation wrappers as seen in our R package, *gwascelltyper*, accessible at: https://github.com/alexandruioanvoda/gwascelltyper

#### ATAC-seq shape-based trend bucker overlap with immune-mediated disease credible set variants

We determined overlap of ATAC-seq shape-based trend bucker peaks with variants from 95% credible sets for various immune-mediated diseases from Farh et al., [33] and Huang et al., 2017 [1]. Credible set variant coordinates were converted from GRCh37 to GRCh38 using the “liftOver” function from the rtracklayer package v.1.54.0, and variants were considered to overlap with a peak if they fell within the start and end of the peak. Credible set variant overlap with read pileup in various activation conditions was plotted using the Gviz package v.1.38.4.

#### Transcription Factor Motif Enrichment

Transcription factor motif enrichment analyses were performed using the “findMotifsGenome” function of the HOMER package v.4.11 [27] with “-size 50 -mask” and all other default parameters. Peaks which were found to be significantly differentially accessible under costimulation conditions as compared to aCD3 stimulation only were compared to a background of all peaks tested in differential accessibility analyses. Peaks which were found to be significantly upregulated under aCD27, aICOS, and aCD6 costimulation conditions as compared to both aCD3 only and aCD3 with aCD28 stimulation were assessed against a background of peaks which were significantly up in these costimulation conditions as compared to aCD3 only conditions. Venn Diagrams were generated using the nVennR package v.0.2.3.

## Supplementary Figures

**Figure S1:**
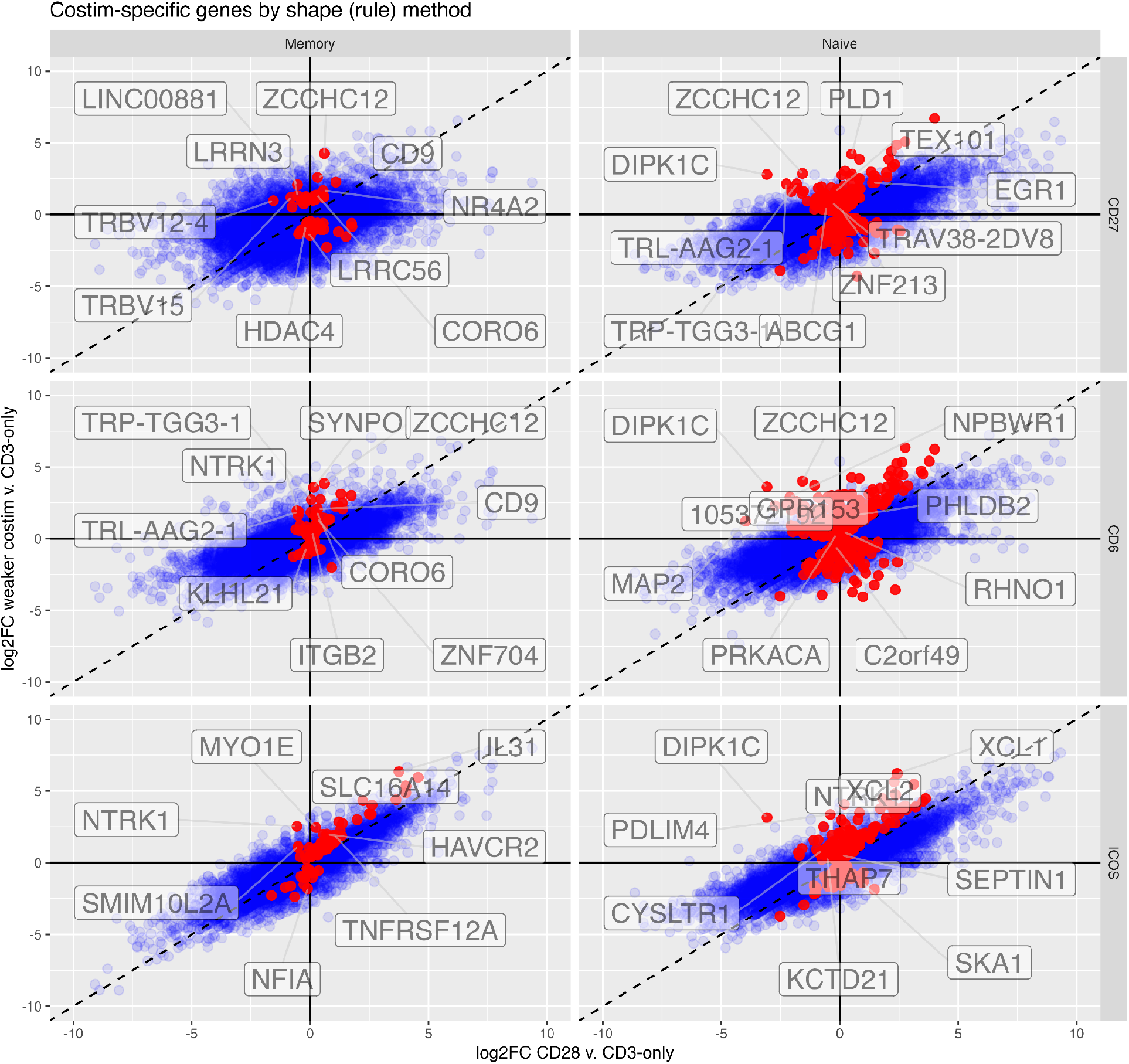
Costimulation-biased genes by the shape method. Gene names are displayed for the top 5 and bottom 5 transcripts ranked by the logFC between the non-canonical costim and CD28. Dashed line is the slope 1 diagonal. The non-canonical significant-bias transcripts are colored in red.

**Figure S2:**
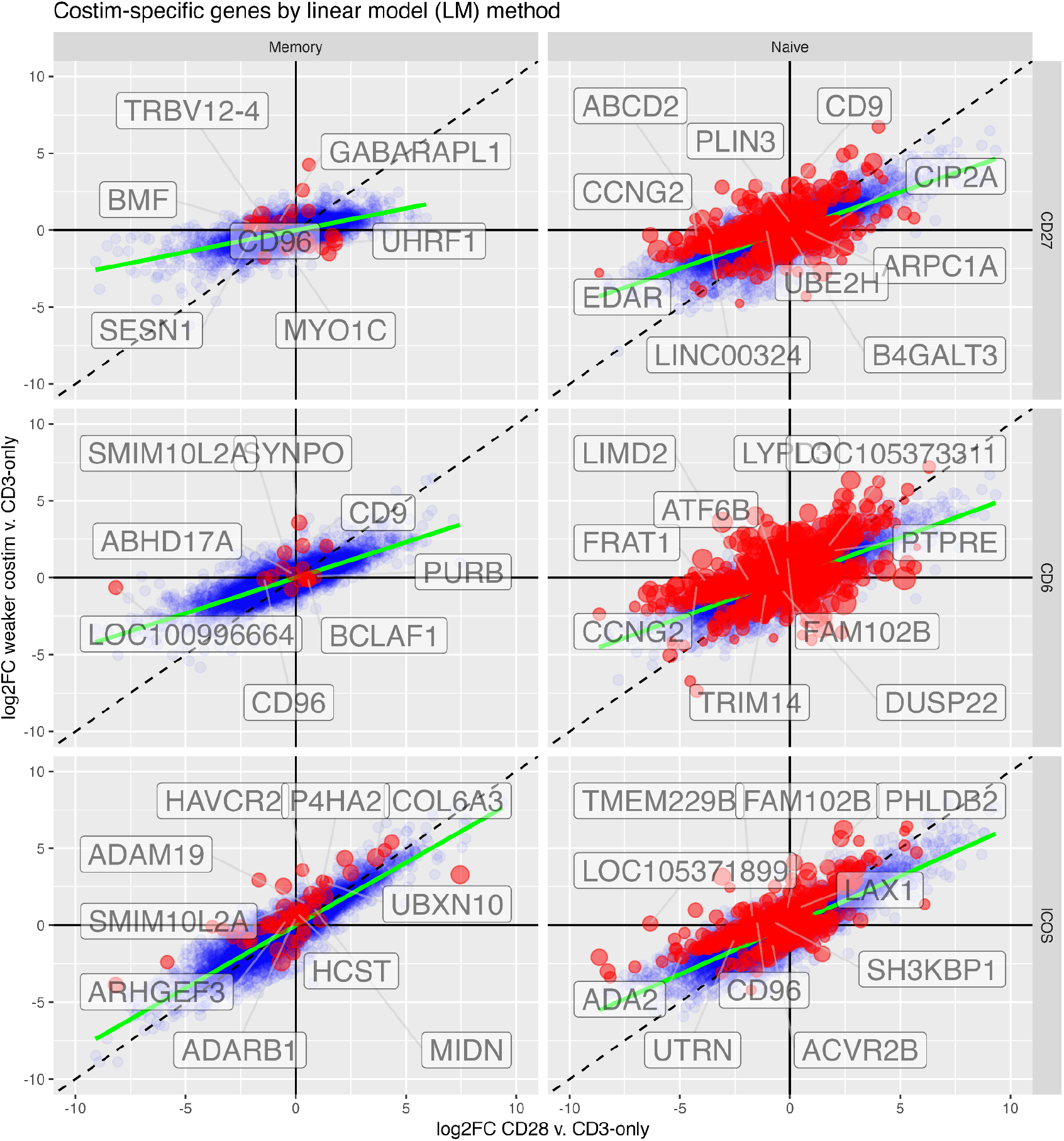
Costimulation-biased genes by the linear model (LM) method. The top 5 and bottom 5 transcripts by Z score are labelled. Dashed line is the slope 1 diagonal, while green line is the linear model fit. Significant transcripts are colored in red, and sized by their absolute Z score.

**Figure S3:**
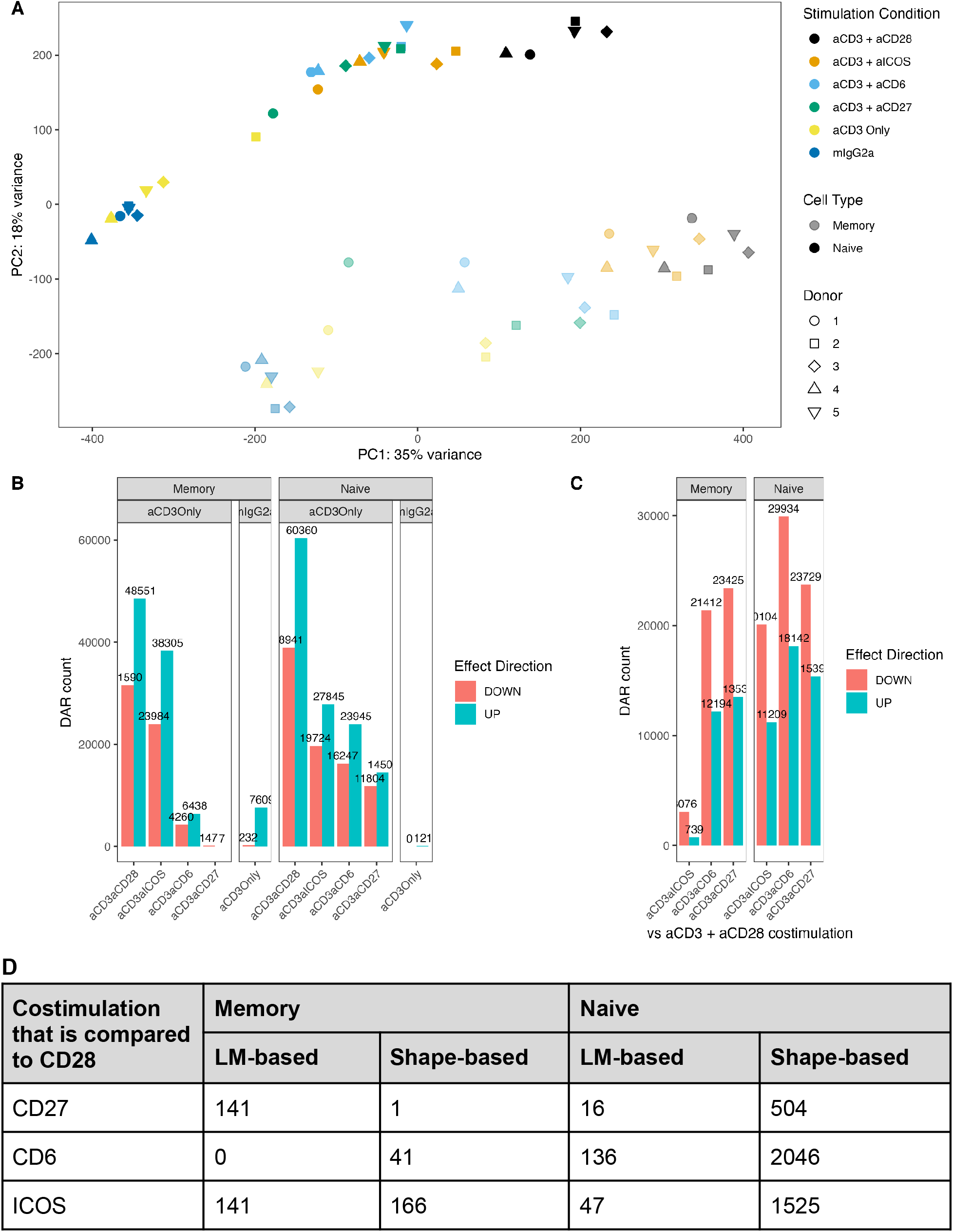
Global changes in chromatin accessibility under different costimulation conditions. **A)** Principal component analysis of ATAC-seq samples, coloured by stimulation condition and shaded by memory/naive subset. **B)** Count of differentially accessible regions (DAR) between costimulation conditions and control conditions, broken down by UP- vs DOWN-regulation, naive/memory subset and control condition (shown in gray boxes above the plot, costimulation conditions are compared to aCD3-stimulated-only controls, and aCD3 is compared to mIgG2a isotype controls). **C)** Count of differentially accessible regions between alternative costimulation conditions (aCD6, aCD27 and aICOS) and the aCD28-costimulated condition, broken down by UP- vs DOWN-regulation and naive/memory subset. **D)** Number of costimulated-biased genes in memory and naive CD4+ T cells under different alternative costimulation conditions compared to CD28 costimulation, estimated using two different methods (LM-based and shape-based).

**Figure S4:**
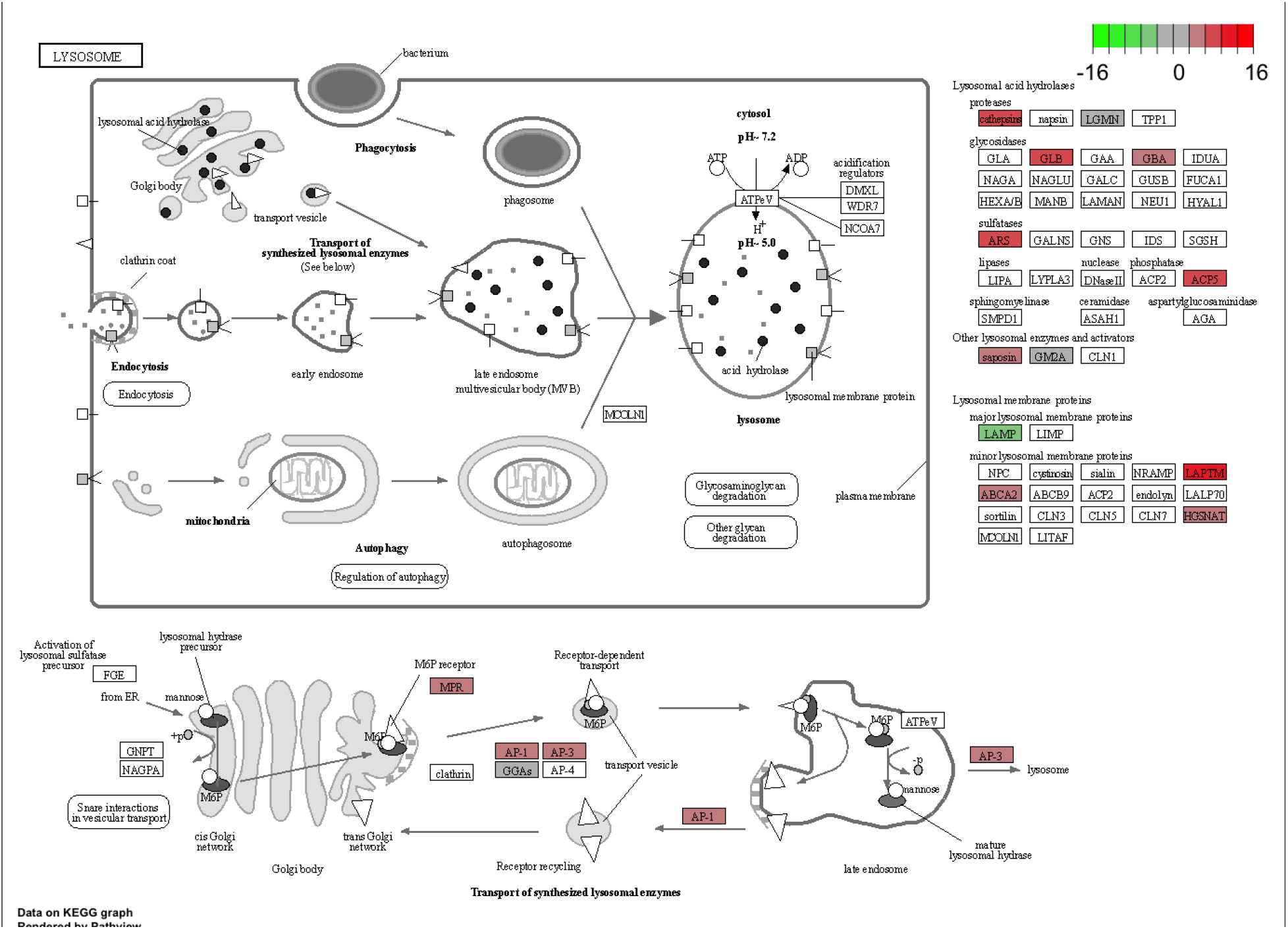
KEGG plot of lysosome genes in naive CD6 v. CD28 costimulation. Positive Z (red) are CD6-upregulated, negative (green) are CD28-upregulated. Figure produced using the R package pathview.

**Figure S5:**
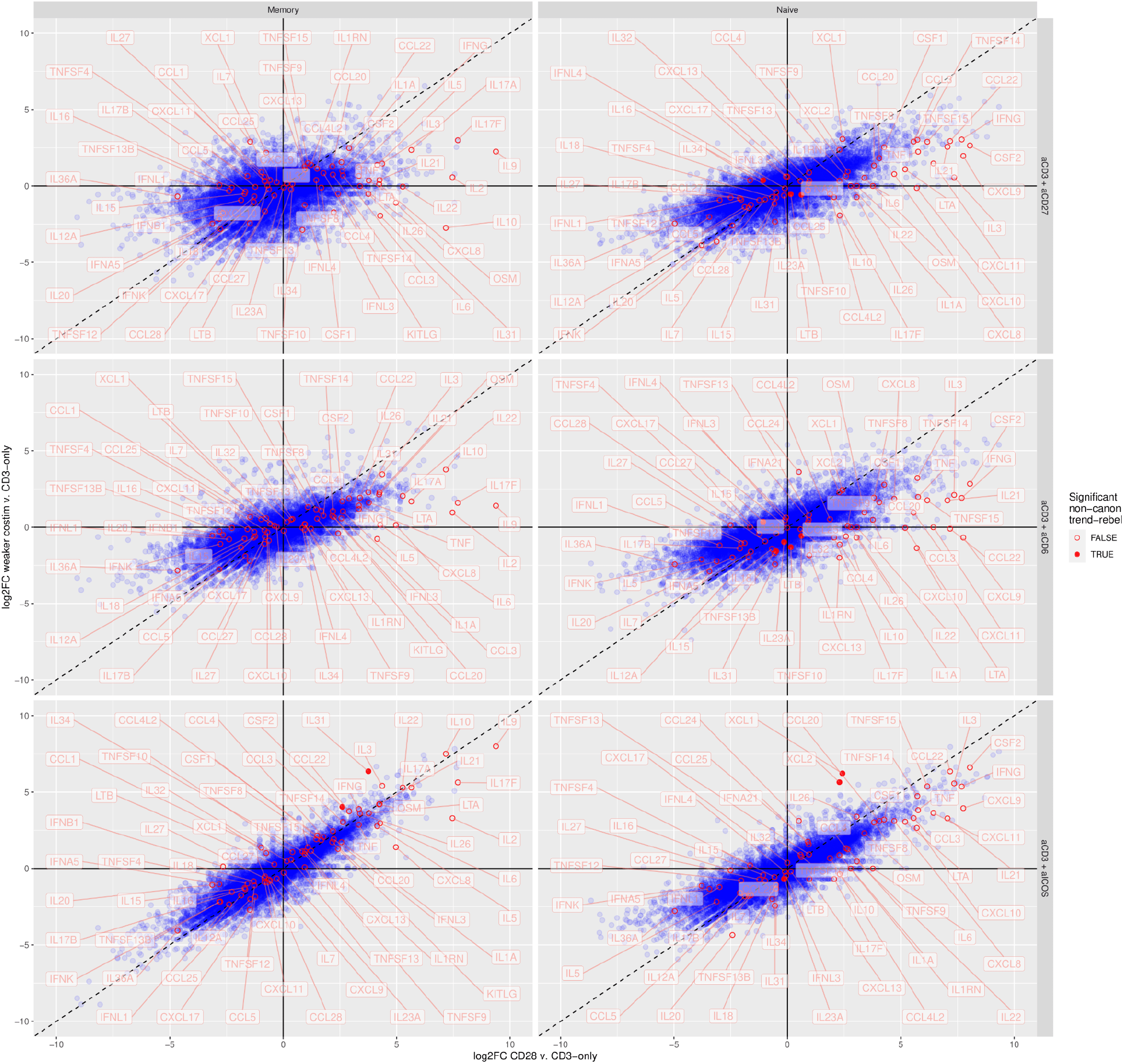
A highlighting of differences in cytokine levels between costimulations. Filled dots represent genes specific to the non-canonical costim based on the shape-method (rule based). The list of cytokines (used for highlighting purposes) was obtained from literature (Santoso et al. 2020).

**Figure S6:**
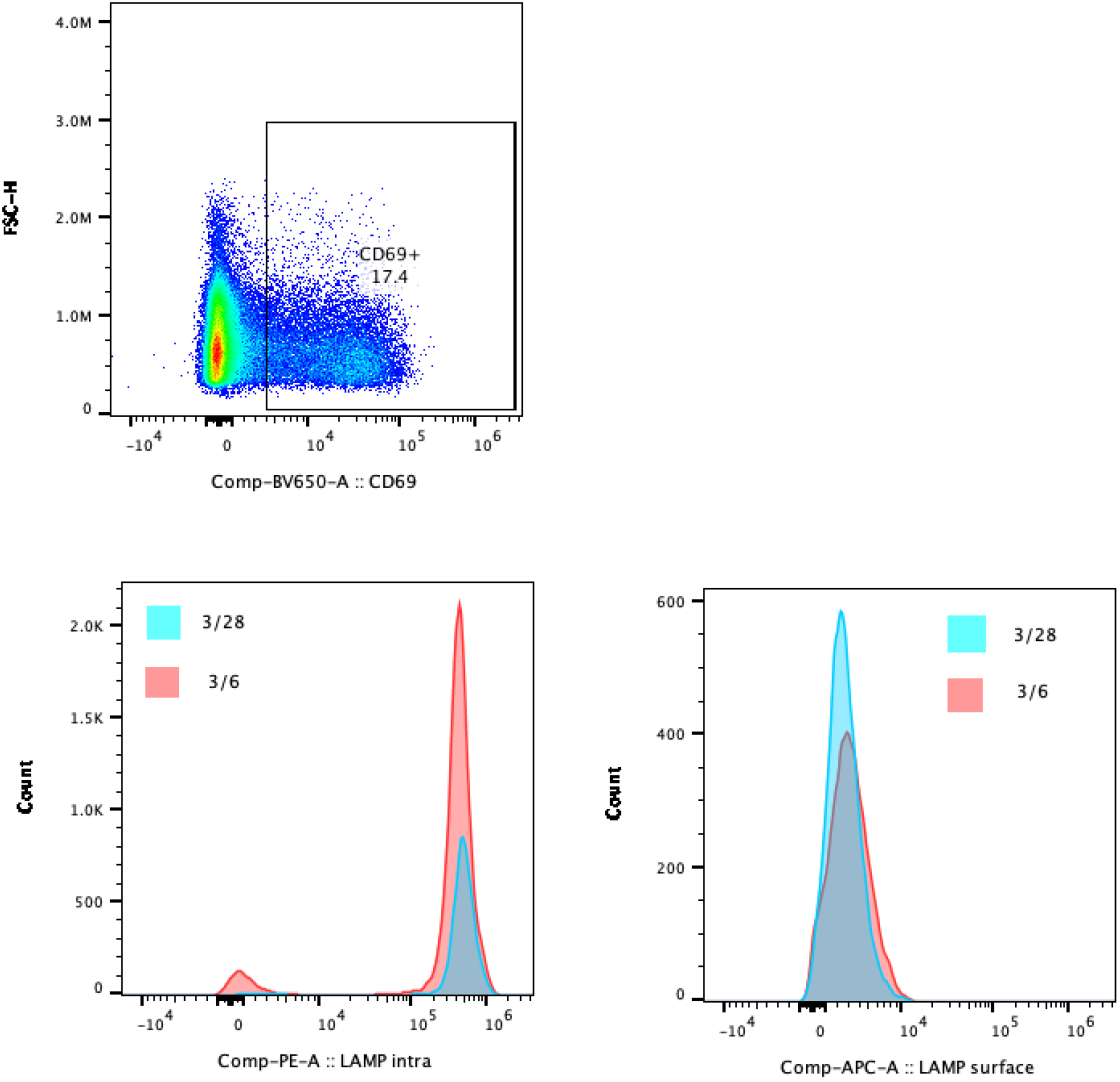
Gating strategy and example expression histograms for measuring LAMP1 expression.

**Figure S7:**
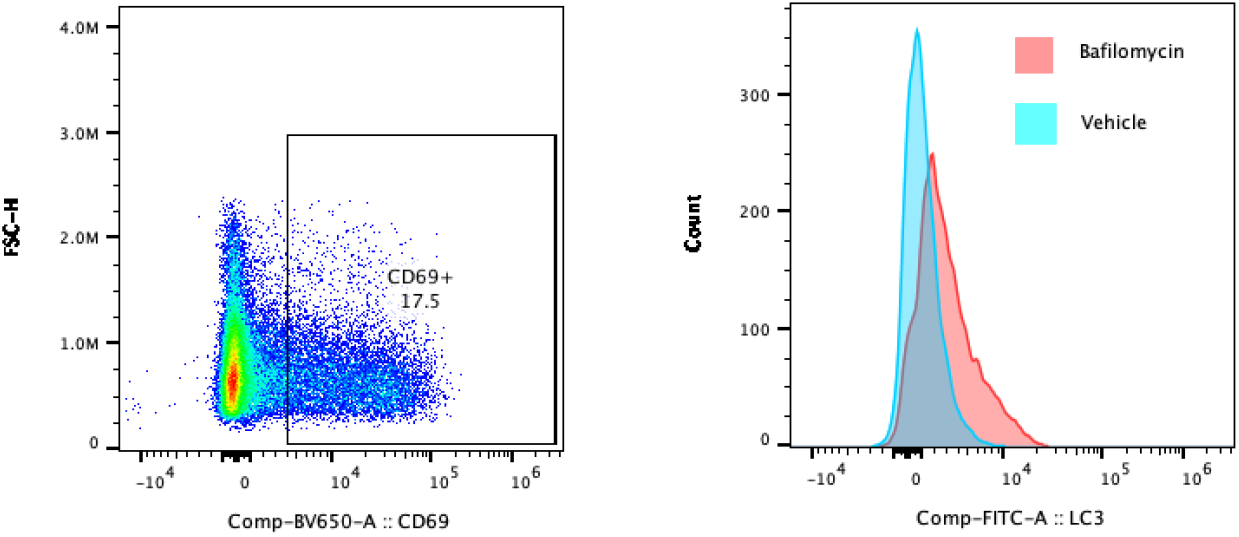
Gating strategy and example expression histograms for measuring autophagic flux expression.

## Supplementary Tables

**Table S1:** Fine-mapped variants for inflammatory disease that lie within costimulation-biased ATAC-seq peaks. Chr=Chromosome, logFC=log fold change, UC=Ulcerative colitis, CD=Crohn’s disease, IBD=inflammatory bowel disease, MS=multiple sclerosis, PBC=primary biliary cirrhosis.

| Region Count | Condition | Chr | Position | rsID | Disease | Fine-mapping posterior | Study | Peak location | logFC costim vs cd3 | logFC CD28 vs CD3-only | logFC costim vs CD28 |
| --- | --- | --- | --- | --- | --- | --- | --- | --- | --- | --- | --- |
| 1 | aCD3aCD6_Naive | chr1 | 2556327 | rs2227313 | UC | 0.00143 | Huang et al | chr1_2555980_2556670 | -0.545 | -0.02 | -0.463 |
| 2 | aCD3aICOS_Naive | chr11 | 96293026 | rs538636 | UC | 0.0647 | Farh et al | chr11_96292738_96293136 | 0.887 | -0.185 | 1.125 |
| 3 | aCD3aCD27_Naive | chr13 | 99384164 | rs9557207 | UC | 0.0189 | Huang et al | chr13_99384021_99384315 | 0.802 | 0.439 | 0.425 |
| 3 | aCD3aCD6_Naive | chr13 | 99384164 | rs9557207 | CD | 0.0189 | Huang et al | chr13_99384021_99384315 | 0.925 | 0.439 | 0.547 |
| 4 | aCD3aICOS_Naive | chr13 | 99434716 | rs7329158 | MS | 0.0303 | Farh et al | chr13_99434243_99434840 | 0.501 | -0.114 | 0.665 |
| 5 | aCD3aCD27_Naive | chr14 | 68285926 | rs8008961 | PBC | 0.0623 | Farh et al | chr14_68285419_68285979 | 0.591 | 0.216 | 0.432 |
| 5 | aCD3aCD6_Memory | chr14 | 68285926 | rs8008961 | PBC | 0.0623 | Farh et al | chr14_68285419_68285979 | 0.533 | 0.235 | 0.351 |
| 5 | aCD3aCD6_Naive | chr14 | 68285926 | rs8008961 | PBC | 0.0623 | Farh et al | chr14_68285419_68285979 | 0.534 | 0.216 | 0.38 |
| 6 | aCD3aICOS_Memory | chr17 | 42362183 | rs744166 | IBD | 0.0635 | Huang et al | chr17_42361869_42362210 | 0.466 | -0.02 | 0.52 |
| 7 | aCD3aCD6_Memory | chr2 | 60844167 | rs4672409 | CD | 0.0168 | Huang et al | chr2_60844140_60844414 | 1.247 | 0.456 | 0.851 |
| 8 | aCD3aICOS_Naive | chr5 | 35877812 | rs10491434 | UC | 0.045 | Farh et al | chr5_35877700_35877979 | 2.277 | 1.354 | 0.971 |
| 8 | aCD3aICOS_Naive | chr5 | 35877739 | rs10491435 | UC | 0.0385 | Farh et al | chr5_35877700_35877979 | 2.277 | 1.354 | 0.971 |
| 9 | aCD3aCD6_Naive | chr8 | 128540606 | rs34841270 | CD | 0.249 | Huang et al | chr8_128540405_128540817 | -0.757 | -0.152 | -0.542 |
| 9 | aCD3aCD6_Naive | chr8 | 128540609 | rs201242438 | CD | 0.0095 | Huang et al | chr8_128540405_128540817 | -0.757 | -0.152 | -0.542 |

